# With great power comes great responsibility: how scientific supervisors shape the wellbeing of early-career researchers

**DOI:** 10.64898/2026.05.05.722947

**Authors:** Xabier Simón Martínez-Goñi, Agustín J. Marín-Peña, Mario Corrochano-Monsalve, Adrián Bozal-Leorri

## Abstract

Scientific supervision is central to the experience of early-career researchers (ECRs), yet its role in shaping wellbeing and retention remains underexamined from the ECR perspective. We analyzed 2,604 anonymous survey responses from predoctoral, postdoctoral and former researchers across 65 countries. Overall, 76% of respondents reported that their supervisor’s attitude had a moderate or severe impact on mental health. Although most entered academia for vocational reasons, negative experiences with supervisors were among the most frequently reported reasons for leaving among former researchers (48%), comparable to job insecurity and financial instability. Harm was most often associated with poor communication, disregard for wellbeing, micromanagement and competitiveness. In contrast, ECRs valued supportive rather than boss-like supervision, regular communication, realistic expectations and respect for personal time. These findings identify supervisory behavior as a major and modifiable determinant or ECRs wellbeing and retention, and highlight the need for stronger institutional accountability, mentor training and funding incentives that recognize mentorship as a core component of research culture.

## INTRODUCTION

Working in academia provides the opportunity to address complex scientific questions, generate new knowledge and contribute to social progress through creativity, collaboration and the training of future generations. Interest in this career path has increased in the last years: between 2014 and 2023, the proportion of people aged 25 to 64 with a doctorate in OECD countries grew by 44%, reaching 1.3% (OECD, 2025). This expansion reflects the growing appeal of pursuing scientific careers and the expectation that a doctoral degree provides meaningful professional opportunities. Yet despite this growth, the experience of doing academic research has become increasingly stressful, shaped by structural pressures and precarious working conditions (Solomon and Du Plessis, 2023). Early-career researchers (ECRs), including PhD candidates and young postdoctoral researchers, face a substantially elevated risk of mental health problems, being over six times more likely than the general population to experience anxiety and depression (Evans et al., 2018; Vigil-Avilés et al., 2024). The personal and professional demands of research are so intense that more than half of those who begin a PhD do not complete it (Lovitts, 2001; Tenorio-Lopes, 2023).

Many challenges only become fully visible once researchers are inside academic structures. Main stressors include scarce academic positions, pressure to publish in high-impact journals, short-term contracts and competitive lab environments (Powel, 2016; Woolston, 2018). In addition, the relationship with the scientific supervisor plays a particularly influential role in shaping the doctoral and postdoctoral experience (Al Makhamreh and Stockley, 2020; Le et al., 2021; Feizi and Elgar, 2023). A supportive and constructive supervisory relationship directly contributes to researchers’ performance, confidence and wellbeing (Sarabipour et al., 2024). High quality mentorship is also associated with persistence in academic careers (Williams et al., 2016). Nevertheless, many ECRs report significant mental health struggles, often linked to inadequate supervision (Gewin, 2012). In the study by Evans et al. (2018), about half of the participants who reported anxiety or depression indicated that their supervisor provided little to no support for their mental wellbeing. Although various authors have proposed concrete strategies to improve academic mentoring (Maestre, 2019; Rillig, 2022), these recommendations appear inconsistently applied, resulting in harmful consequences for young researchers.

Much of the discussion around supervision and mentoring continues to be shaped by the perspectives of principal investigators or by institutional and country-skewed datasets (Tenenbaum et al., 2001; Levecque et al., 2017; Tenorio-Lopes, 2023; Sarabipour et al., 2023; 2024), leaving limited space for the voices and lived experiences of ECRs. Motivated by our own observations of how ECRs’ expectations and needs are frequently overlooked in everyday academic practice, we sought to articulate these perspectives in a more systematic way. This article, written by ECRs, aims to help correct that imbalance by centering the ECRs’ perspective as a source of systematic evidence rather than anecdote. To do so, we collected 2,604 anonymous responses from predoctoral, postdoctoral and former researchers across 65 countries (Fig. 1), generating one of the broadest international datasets focused specifically on how supervision is experienced from below. By foregrounding these accounts, our work identifies recurrent supervisory behaviors, quantifies their impact on wellbeing and career trajectories and outlines evidence-based priorities for improving mentorship. Rather than describing isolated problems, the patterns emerging from this diverse global sample reveal structural issues, entrenched power dynamics and clear opportunities for institutional change.

**Fig. 1.**
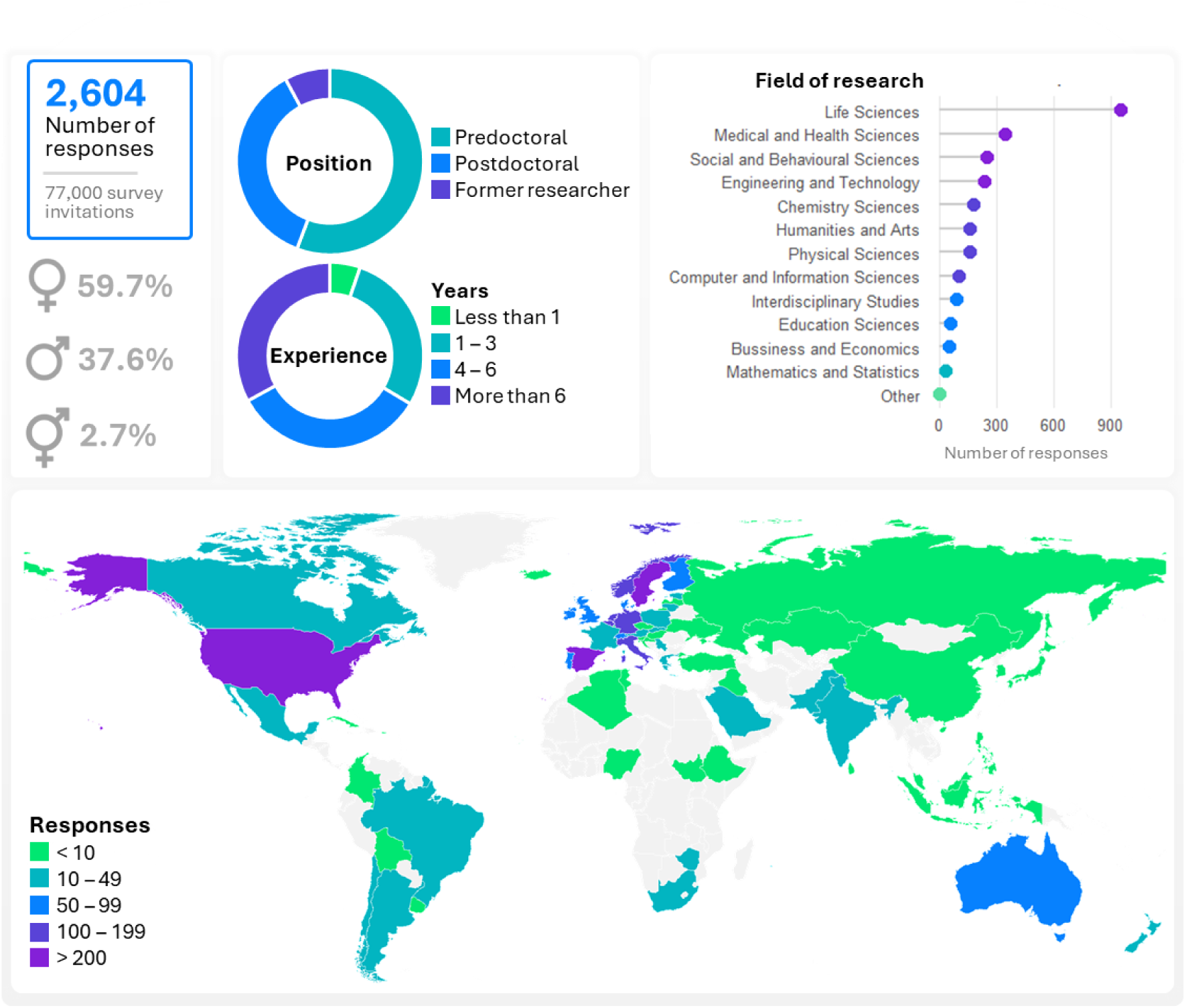
Respondent overview and sample composition. Panels summarize the survey sample, including gender distribution (female, male, and non-binary/other), career position or stage, and years of research experience. The dot plot shows the number of responses by field of research, whereas the map shows the geographic distribution of respondents (note that this indicates the country where the respondent was working at the time of completing the survey, not their nationality).

## METHODS

### Survey design and dissemination

We collected responses through an online survey administered via Google Forms between December 2024 and August 2025. Before beginning the survey, all participants were provided with a clear statement regarding the objectives of the research. They were informed that their participation was entirely voluntary and that they could withdraw from the survey at any point without providing a reason. Furthermore, we assured all respondents that the survey was strictly anonymous, and that no personal identifiable information was collected. Consequently, informed consent was obtained from all participants prior to their involvement in the study.

The questionnaire consisted of closed-ended and multiple-choice items addressing supervisory practices, communication patterns, workload management, career development, and overall wellbeing. To ensure broad international dissemination, we contacted more than 77,000 researchers through institutional mailing lists of universities worldwide and promoted the survey through the authors’ professional networks on X, Bluesky, LinkedIn, and academic forums. Participation was voluntary and no compensation was provided. Recruitment was conducted through broad, non-targeted dissemination channels, without any intention to focus on or overrepresent specific demographic groups, research fields or countries.

### Sample description

Our sample included 2,604 responses from predoctoral (56%), postdoctoral (36%), and former researchers (8%), enrolled in more than 13 different fields of research, the most common Life Sciences (37%), followed by Medical and Health (13%) and Social and Behavioral Sciences (10%) (Fig. 1). Respondents were working in 65 different countries at the time they completed the survey, with the largest shares from Spain (18%), Sweden (11%), and the United States of America (8%). Funding status was fellowships/grants (45%), group-funded positions (45%) and other sources (10%). Participants self-described as female (60%), male (37%), and non-binary/others (3%), and self-identified as White (75%), Asian (10%), Latino (6%), Black (3%), mixed (2%), and others (3%) (Supplementary Fig. S1).

Participants were asked to answer the survey questions with a particular supervisor (of their choice) in mind. The participants chose their supervisor during the predoctoral stage (65%), the early postdoctoral (19%), and the experienced postdoctoral stage (16%). While the sample includes mid-career researchers (experienced postdoctoral stage), all respondents are described as early-career researchers (ECRs) to maintain clarity, as the vast majority (84%) fall into this category. Supervisors were identified as male (61%), female (38%), and non-binary/others (1%), and described as White (87%), Asian (6%), Latino (3%), mixed (2%), Black (1%), and others (2%). Given that approximately 80% of responses originated from Europe, the dataset reflects a strong regional concentration, which should be taken into account when considering the global generalizability of the findings.

## RESULTS AND DISCUSSION

### From vocation to exit: reasons for leaving academia

Academic careers are often portrayed as a natural extension of scientific vocation, a path sustained by intellectual curiosity, long-term commitment, and the expectation of meaningful contribution. Yet, despite this strong initial motivation, the academic trajectory is far from linear, and many researchers ultimately decide to leave it behind (Naddaf, et al., 2024). Although most respondents reported vocational motivation to enter academia (80.5%) (Fig. 2a), departures remain common. Among former researchers (8%), negative experiences with their supervisor emerged as one of the most frequently cited reasons for leaving (48%) (Fig. 2b), highlighting the central role that supervision plays in retention and overall wellbeing. This proportion was comparable or higher than other structural stressors, such as the lack of stable job opportunities (52%), low salaries or limited financial stability (50%), and concerns related to work-life balance (47%), as well as high pressure or stress (43%). In contrast, only 35% indicated that they left academia primarily to pursue other professional goals, underscoring that voluntary career redirection was far less common than departures linked to adverse working conditions or relational dynamics.

**Fig. 2.**
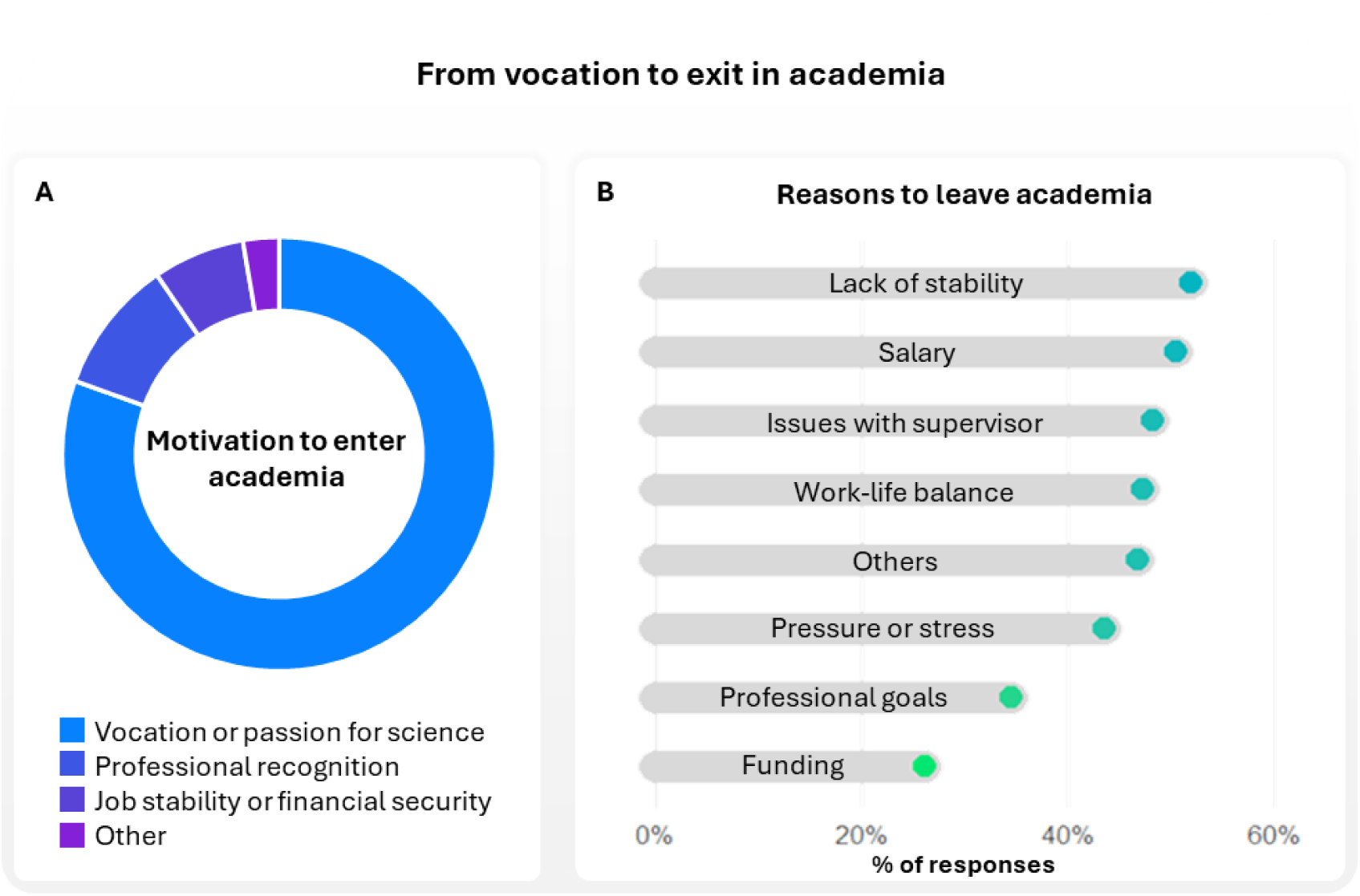
From vocation to exit. **A)** Reported primary motivation for entering academia. **B)** Distribution of reported reasons for leaving academia (respondents could select more than one option).

These patterns portray a career landscape that can extract a substantial mental toll and sharpen the stakes of everyday supervision. While supervisors cannot always address systemic issues such as funding volatility or job precarity, respondents’ accounts indicate that they nevertheless function as a decisive local lever for retention, wellbeing, and progress. A supportive supervisor may buffer structural pressures through predictable, bidirectional communication, calibrated expectations, respect for time outside work, and active engagement in professional development; conversely, poor supervision may amplify the very stressors that precipitate exit (Levecque et al., 2017). Below, we examine the possible reasons why relationships with supervisors are among the main factors driving scientists to leave academic careers. We identify key harmful patterns and highlight good practices that could help retain scientific talent in academia.

### What hurts: Mental health impact of supervisory misconduct

Overall, 76% of respondents reported that their supervisor’s attitude had a moderate (31%) or significant (45%) impact on their mental health (Fig. 3a). The behaviors more often reported as harming morale and productivity were dismissive or disrespectful communication (selected by 52%), little or no feedback on performance (50%), disregard for personal circumstances and well-being (44%), micromanagement (40.5%), and promoting competitiveness among team members (37%) (Fig. 3b). This emphasizes that the most damaging supervisory actions stem from a fundamental lack of respect for the ECR’s professional and personal boundaries, poor communication, and the absence of constructive professional development support. This pattern is consistent with previous studies showing that the quality of the supervisor-ECR relationship is one of the strongest predictors of anxiety and depression among doctoral students (Jackman et al., 2022; Jones-White et al., 2022).

**Fig. 3.**
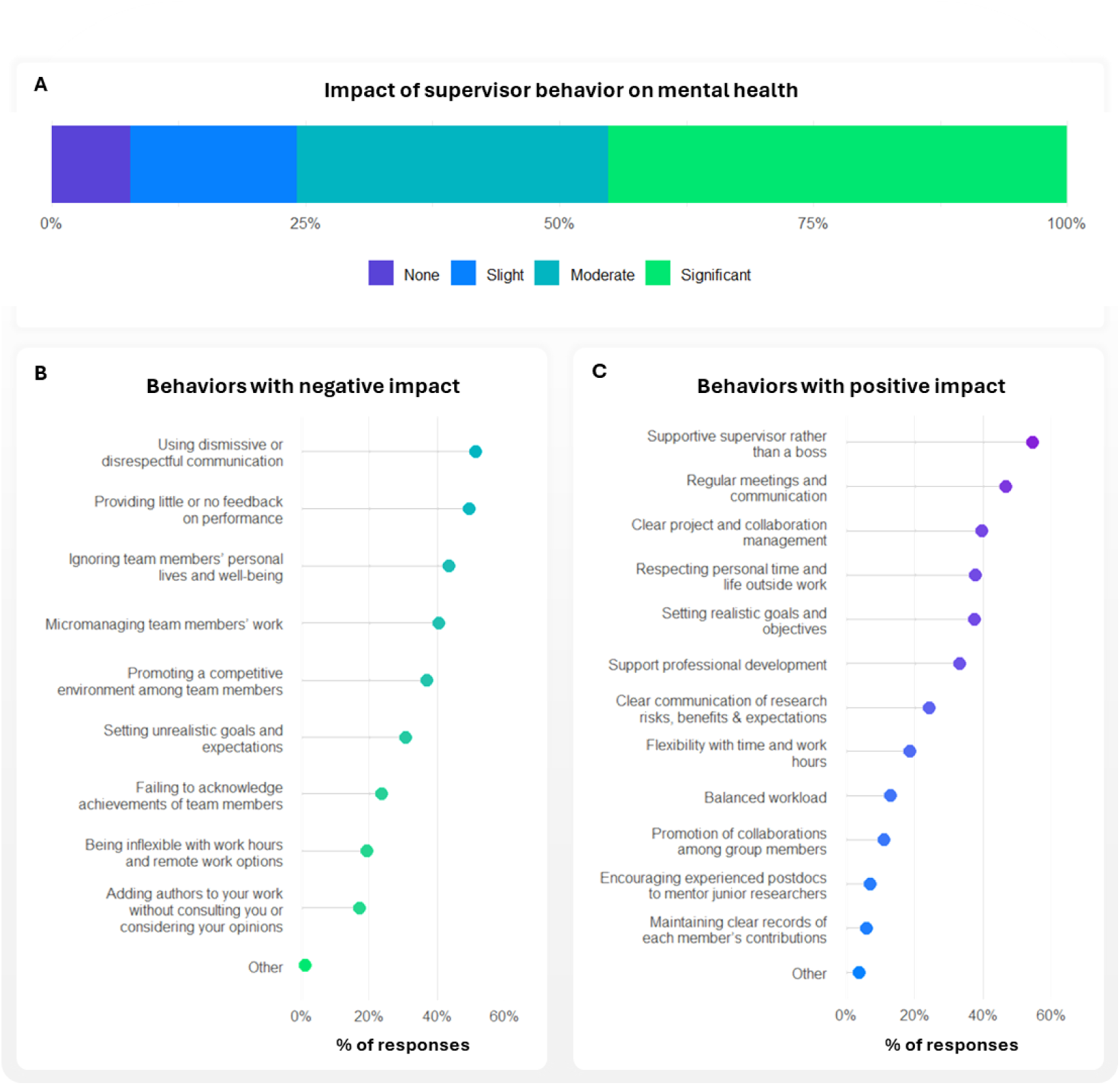
What hurts, what helps: mental health in supervisor - ECR relations. Perceived impact of supervisor behavior on mental health and frequency of particular behaviors. **A)** Overall perceived impact of supervisors’ behavior on respondents’ mental health. **B)** Percentage of respondents reporting specific supervisor behaviors with a negative impact. **C)** Percentage of respondents reporting specific supervisor behaviors with a positive impact. Respondents selected their top three behaviors each category (positive or negative).

Additionally, we asked the participants to identify particular negative experiences with their supervisors during their career (Figs. 4 and Supplementary Fig. S2). These experiences clustered into five prevalent bands. High-frequency issues (reported by ∼30% to ∼40% of respondents) included disorganization (38%) and lack of communication (37%), and relational instability or boundary breaches: abrupt attitude changes (33%), lack of support (31%), lack of empathy (31%), and contact outside working hours (30%). These results indicate that the majority of negative experiences reported by ECRs in the research environments result from deficiencies in supervisory management (disorganization, lack of communication, intrusion on personal time) as well as relational shortcomings. In the same manner, these high-frequency issues suggest failures in establishing clear professional boundaries and effective management practices within supervisory roles.

**Fig. 4.**
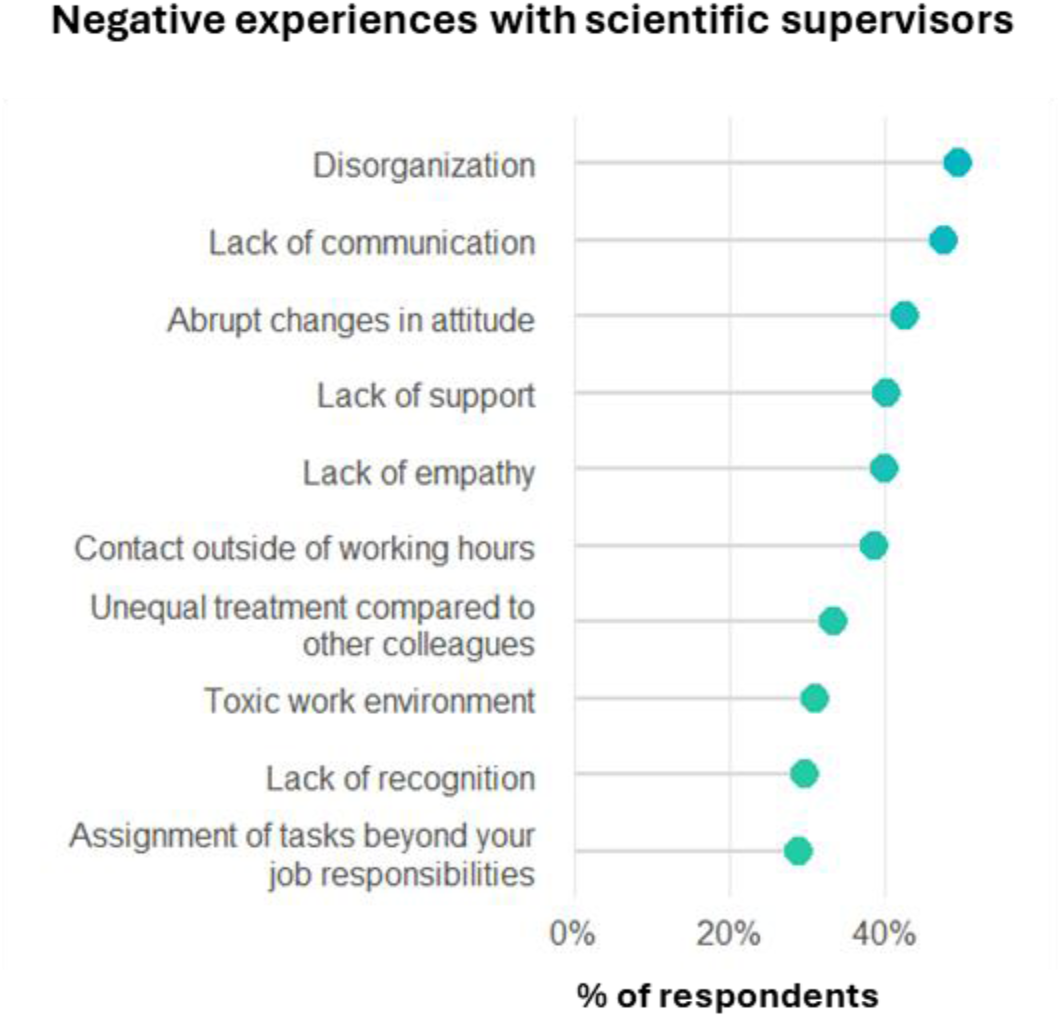
Reported negative experiences with scientific supervisors. Percentage of respondents reporting each situation (10 most frequently reported situations are shown; the remaining are shown in Supplementary Fig. S2. Participants were asked to select all situations they had experienced.

This finding is not particularly surprising, given that advancement to a supervisory role typically relies on securing competitive funding and resources, a process that rarely includes formal preparation in leadership or mentorship skills (Lacey et al., 2025). Intermediate issues (reported by ∼20% to ∼30% of respondents) included unequal treatment (26%), toxic work environment (24%), lack of recognition (23%), tasks beyond responsibilities (23%), lack of resources (22%), criticism/exclusion (21.5%) and feeling of exclusion (21.5%). Thus, beyond the interpersonal and organizational shortcomings, the second block highlights structural challenges in how ECRs’ professional roles are defined, valued, and evaluated within research groups by their supervisors. Such issues point to systemic weaknesses in management practices and institutional oversight, which may perpetuate inequities and hinder the development of supportive, transparent supervisory relationships. Less common but consequential issues (13% - 20% prevalence) included inappropriate comments (20%), micromanagement (19%), expectations of loyalty/gratitude (16%-18%), interference in personal life (15%), and breaches of agreements (13%). Although these behaviors did not reach the high prevalence rates of organizational failures, the fact that up to one-fifth of respondents experienced such experiences confirms that many supervisors routinely fail to respect the professional autonomy and personal space of their ECRs. Severe misconduct was reported by a minority (< 10% prevalence) including verbal violence (9%), threats (7%), and sexual harassment/physical violence (< 1%), but remains critical given its potential harm. Specifically, the fact that verbal violence and threats combine for over 15% indicates that a substantial minority of ECRs are subjected to emotionally damaging and professionally unsafe supervisory conduct. While less frequent than managerial and organizational failures, these findings underscore that serious forms of supervisory misconduct persist within academia and demand targeted institutional interventions, clear accountability mechanisms, and comprehensive mentor training programs (Hund et al., 2018; Busch et al., 2024).

Given that many of the negative experiences described above reflect hierarchical supervisory practices, blurred professional boundaries, and forms of dependence that can undermine ECRs’ autonomy, we next asked whether their funding source might also be perceived as relevant to supervisory dynamics. Interestingly, most ECRs (68%) considered that their supervisor’s attitude varied depending on the funding source, either somewhat (38%) or significantly (30%), whereas 19% reported no difference (Supplementary Fig. S2). This finding suggests that financial dependence may further modulate how supervisory authority is exercised and experienced, opening an important avenue for future research.

### Supervision and wellbeing: Patterns of support

Notably, 22% of participants reported no negative situations with their supervisors. While the specific experiences of these particular ECRs with their supervisors are unknown, it can be reasonably inferred that their supervisors embody many of the positive characteristics identified as key by the general respondent pool in which respondents selected the top three characteristics of a supportive supervisor (Fig. 3c). The most frequently selected were being supportive rather than boss-like (54.5%), regular meetings/open communication (47%), clear project and collaboration management (40%), respect for personal time (38%), and realistic goals/objectives (38%). This high-frequency block highlights that researchers primarily value empathy, clear organization, and healthy boundaries as the core requirements of good supervision. Supporting ECRs’ professional development was selected by 33%, while clarity about research risks/expectations (24%) and flexibility with time and work hours (18.5%) were also valued. Therefore, it becomes evident that maintaining open communication, holding regular meetings, establishing clear and realistic objectives, and respecting life outside work are essential elements of a healthy supervisory relationship (Cornejo-Araya et al., 2024). Yet, the most critical factor appears to be having a supportive supervisor rather than a boss-like one who views ECRs merely as producers of scientific outputs such as data and publications. Indeed, many other positive supervisory traits, such as avoiding work overload, setting attainable goals, respecting personal boundaries, and fostering mutual respect, naturally stem from being supportive and empathetic. Treating ECRs as colleagues at an earlier stage of the same academic journey, rather than as production units, may lie at the core of effective mentorship (Powell et al., 2016).

### Communication as a core element of supervision

Regular meetings and open communication were consistently identified as core to a healthy supervisory relationship. Nearly all ECRs (∼90%) rated meetings as important or extremely important (Fig. 5a). Most judged the number of meetings as sufficient (62.5%), even though 31% considered the number of meetings insufficient (Fig. 5b). The high perceived value is further confirmed by the quality assessment, where the majority of respondents (75%) found meetings useful (51%) or extremely useful (24%) (Fig. 5c). Nonetheless, still 25% reported useless (18%) or even a complete waste of time (7%). Importantly, “more” was not necessarily “better”. Among 7% reporting too many meetings, perceived utility dropped sharply: useful/extremely useful fell to 38% (vs 75% overall), and poor-quality meetings rose to 62% (41.5% useless; 20.5% waste of time). These results support prioritizing predictable rhythms and high-quality interactions over meeting volume.

**Fig. 5.**
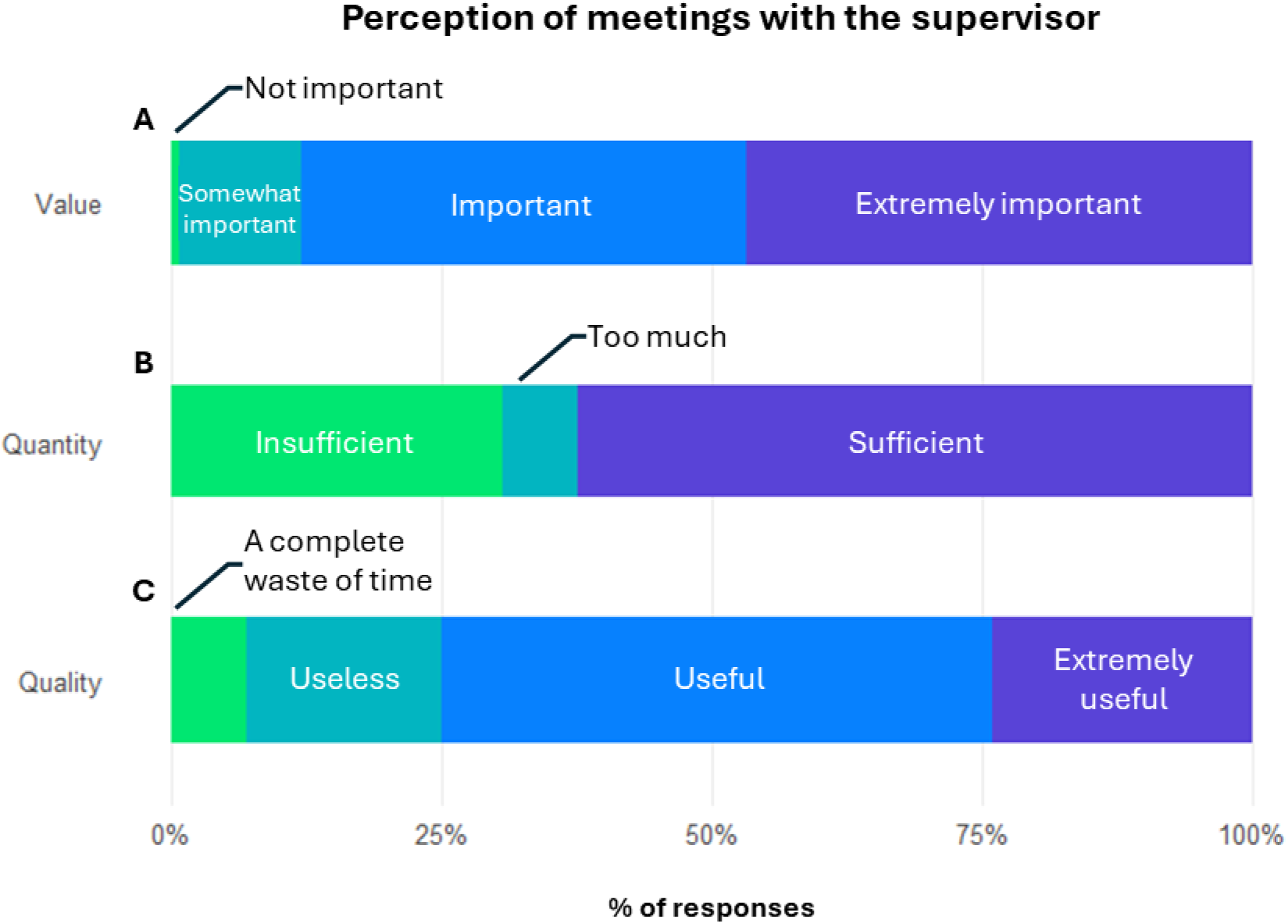
Communication as a core element of supervision. Perceived value, quantity, and quality of supervisory meetings. Bars show respondents’ ratings of **A)** the importance of regular meetings, **B)** their frequency, and **C)** the perceived usefulness of meetings.

Crucially, these meetings also serve as a key space for involving ECRs in research related decisions, reinforcing their sense of collaboration, shared responsibility and professional growth within the team (Sarabipour et al., 2023; 2024). This connection is strongly reflected in participants’ responses: fewer than 1% considered inclusion in decision making unimportant, while 92% rated it as important (50%) or extremely important (42%) (Supplementary Fig. S4). Such results indicate that structured communication is not only valued for monitoring scientific progress but also for enabling ECRs to feel meaningfully engaged in the research process and treated as early stage colleagues rather than passive executors.

Taken together, these findings underscore that predictable, high-quality meetings and shared decision making operate jointly as central pillars of effective supervision, shaping ECRs’ autonomy, motivation and perception of being part of a genuine collaborative effort (Sarabipour et al., 2023; 2024).

### Time management, workload and boundaries

While communication and inclusion define the working relationship, ECRs stressed a core requirement for a model supervisor: the recognition of the human factor in aspects such as workload, empathy for personal circumstances, and respect for time outside workThis emphasis resonates with academic discussions that identify respect for personal boundaries, empathy, and attention to a student’s life circumstances as core contributors to doctoral wellbeing and positive supervisory relationships (Rillig, 2022; Cornejo-Araya et al., 2024), but this is particularly relevant given the prevalence of work-life imbalance. Only 44% reported working within scheduled hours while maintaining a healthy personal balance (Fig. 6a). In contrast, the majority (56%) reported working beyond normal hours (35%) or making constant personal sacrifices (21%). Despite this, most respondents reported being generally satisfied (55%) or very satisfied (13%) with their current work approach (Fig. 6b). However, the remaining 31% who reported absolute or slight dissatisfaction underscored a persistent failure to manage work-life balance effectively. Among dissatisfied respondents, working hours beyond scheduled increased to 42% (vs. 35% overall), and constant personal sacrifices increased to 44% (vs. 21%). Among fully dissatisfied ECRs, constant personal sacrifices rose to 62%, indicating that perceived personal cost, more than long hours alone, tracks strongly with dissatisfaction.

**Fig. 6.**
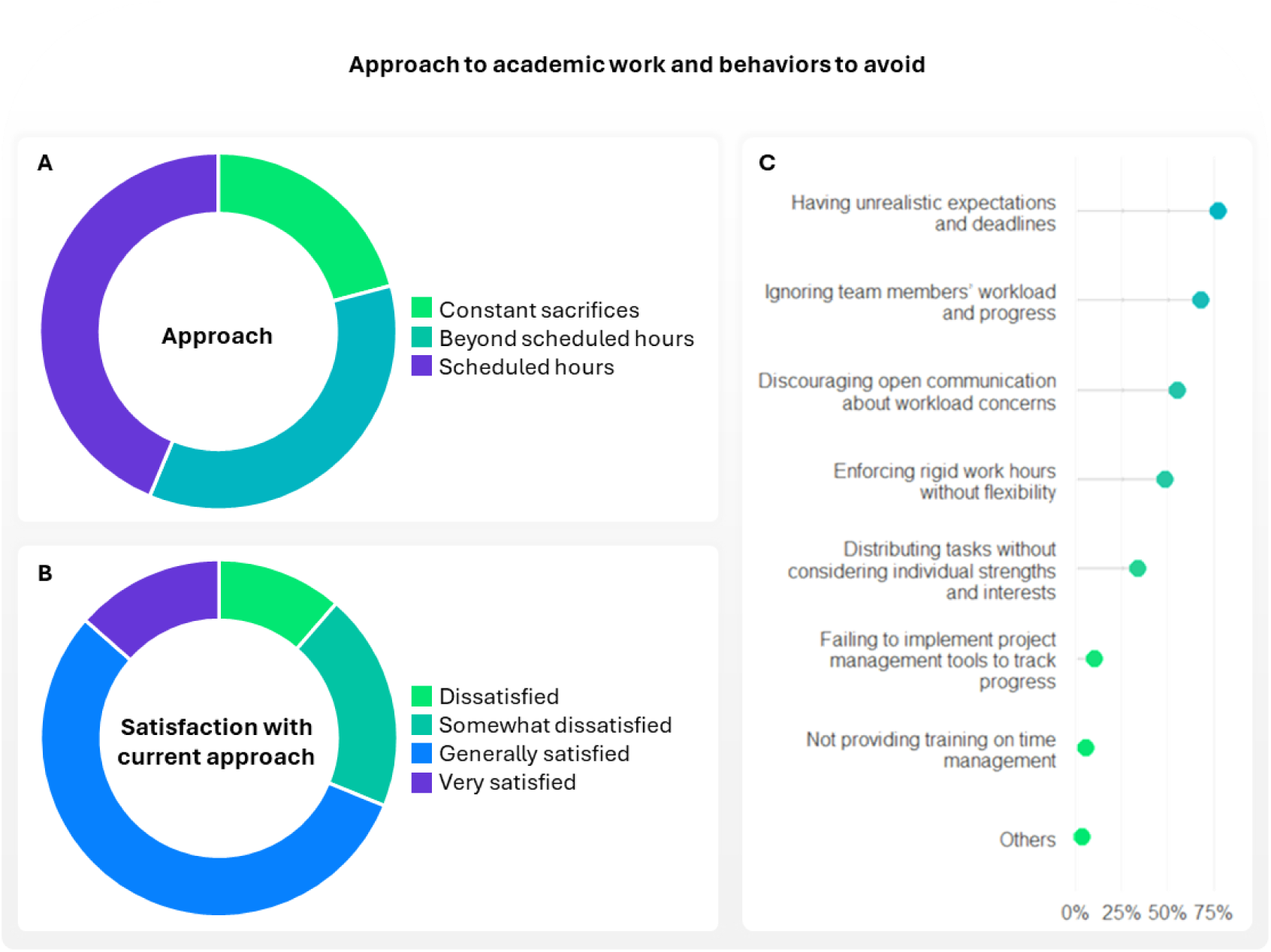
Time management, workload and boundaries. **A)** Distribution of respondents’ usual approach to academic work. **B)** Satisfaction with current approach. **C)** Specific practices that respondents believe supervisors should avoid to maintain balanced workloads (respondents were asked to pick three behaviors).

These results suggest that excessive and overwhelming workload is among the primary drivers of negative environments and poor ECR well-being, severely impacting their relationship towards academic work (Busch et al., 2024), This pattern points to a clear association: while working beyond normal hours is common across the entire sample, the experience of constant personal sacrifice, a core indicator of unhealthy work-life balance, is more than tripled among the most dissatisfied ECRs (21% general vs. 62% fully dissatisfied). This highlights the critical importance of effective workload management in maintaining a healthy and sustainable research environment. This persistent failure is attributable to complex factors, which may include the demanding academic system dynamics (“publish or perish”), lack of institutional action, and ineffective supervisory management (Powel, 2016; Woolston, 2018).

When asked which strategies scientific supervisors should take to maintain balanced workloads, ECRs overwhelmingly selected realistic expectations and deadlines (82%) (Fig. 6c) encouraging open communication about workload (60%), and regular check-ins on workload/progress (53.5%). These findings position supervisors as key local levers for workload sustainability, while institutional policy is needed to mitigate system-level pressure. Given the supervisor’s role as the direct manager of the team and mentor, they serve as the key facilitator for ensuring effective work-life balance and recognition of the human factor for their team members. Institutional policy must then support this effort by mitigating system-level pressure. If these actions are not consistently implemented at both supervisory and institutional levels, the detrimental pressures of the academic system will remain unchanged and continue to erode ECR wellbeing (NASEM, 2019).

### After-hours group leisure activities

A delicate test of how supervisors manage time boundaries and respect life outside work concerns after-hours group leisure activities. Although social interactions outside formal work settings are often assumed to strengthen group cohesion and foster a positive laboratory climate, their effects on ECRs can be more complex and depend on personal circumstances, workload, and the quality of the supervisory relationship. Nearly half of supervisors (45%) were reported to organize after-hours leisure activities (Supplementary Fig. S5), indicating that such practices are common in many research environments. Among respondents whose supervisors organized these activities, 78% enjoyed them, whereas 22% felt obligated to participate, highlighting that the same activity can be experienced as either beneficial or burdensome depending on individual circumstances. Indeed, what some ECRs find energizing, others may perceive as exclusionary, impractical, or unwelcome because of family responsibilities, financial constraints, health, social anxiety, or a desire to maintain clearer boundaries between work and personal life. Consistent with this, 33% of ECRs reported that they either feel obligated or would not like to attend such events, suggesting that a potentially positive practice can also become a source of pressure. At the same time, among respondents whose supervisors did not arrange leisure activities outside work, a substantial proportion (59%) said they would have enjoyed them, pointing to a missed opportunity to foster cohesion. Together, these findings suggest that after-hours social activities can be valuable, but only if participation is clearly voluntary and declining carries no penalty. Open communication is therefore essential, and such activities should be complemented by equivalent opportunities for relationship-building during working hours, so that social engagement supports team cohesion without compromising autonomy or personal well-being.

### Shaping the ideal supervisor

Across our findings, effective mentorship emerges as a combination of empathy, relational competence, clear management, and respect for the ECR’s autonomy. Rather than pointing to a single desirable trait, respondents’ answers suggest that good mentorship depends on several closely connected dimensions that shape both day-to-day working conditions and long-term professional development.

First, respondents clearly valued supervisors who are supportive rather than boss-like. This preference suggests that ECRs do not simply expect technical guidance, but a supervisory approach grounded in respect, professional recognition and a genuine commitment to their development. In practice, this implies treating ECRs as colleagues at an earlier stage of the same academic journey, rather than primarily as producers of scientific output. Such an approach requires prioritizing proactive mentorship and guidance and, obviously, avoiding disrespectful communication or verbal abuse/threats, which were among the most harmful experiences reported in our dataset.

Second, open, regular, and respectful communication appears to be a central pillar of effective supervision. Regular meetings, useful feedback, workload calibration, and clear project management, help create clarity around expectations, strengthen collaboration and support ECRs’ sense of agency within the research process. These practices are especially important considering the high frequency of reported communication failures and lack of guidance (Sarabipour et al., 2023; 2024).

Third, respondents highlighted the importance of realistic goals and active workload management. Unrealistic expectations were identified as one of the most problematic supervisory behaviors, and excessive workload was closely associated with dissatisfaction and personal sacrifice. Effective supervision therefore requires not only scientific ambition, but also the ability to calibrate expectations, monitor workload and foster a collaborative rather than a competitive environment.

Finally, good supervision also depends on respect for personal boundaries and individual autonomy. Respondents consistently emphasized the need for supervisors to recognize the human factor, including personal circumstances, wellbeing and life outside work. Allowing flexibility with time, work without enforcing rigid hours, and avoid interfering in personal life or creating expectations of loyalty or gratitude that prioritize work over personal life, appear to be important conditions for maintaining a healthy and sustainable supervisory relationship.

Taken together, these directives are best understood not as isolated supervisory behaviors, but as interconnected expressions of a broader underlying quality: empathy. When supervisors approach mentorship with genuine empathy and respect for the ECR’s position, the remaining positive practices, such as being supportive, setting realistic goals, and respecting personal boundaries, arise naturally. At the same time, effective mentorship cannot be entirely standardized, as it requires adaptation to the specific needs, background and career stage of each ECR.

### Systemic solutions for improving supervisory practices

#### The role of Research Institutions

Improving supervisory practices cannot depend solely on the goodwill or individual competence of individual supervisors. The prevalence of negative experiences including disorganization, poor communication, lack of support and, in some cases, severe misconduct points to structural weakness that require institutional intervention. Universities and research centers should therefore shift their focus from treating supervisors merely as funding gatherers and publishing entities to recognizing them as essential managers and mentors whose practices directly shape the wellbeing, development and retention of early-career researchers (Pizzolato and Dierickx, 2023; Rahal et al., 2023).

One important implication is that mentorship should be treated as a professional competency requiring formal preparation and continued oversight. Institutions could support this by implementing mandatory training in leadership, mentorship, conflict management and responsible supervision, complemented by refresher courses and regular anonymous ECR evaluations reviewed by independent committees.

At the same time, institutions should create stronger incentives for good supervision, which should be formally recognized in performance reviews and promotion processes. Public recognition, mentorship awards, financial bonuses, or temporary reductions in teaching or administrative duties could also help to highlight that supportive supervision is a valued component of academic excellence. Conversely, repeated evidence of harmful supervisory practices should trigger proportionate responses, ranging from remedial training and improvement plans to formal review procedures, negative punctuation for internal promotions and, where necessary, temporary restrictions on recruiting new researchers or accessing certain internal resources.

#### The role of Funding Bodies

To promote a culture of excellence in supervisory practices beyond institutional walls, funding bodies have an important role in promoting better mentorship. When grant systems prioritize scientific output almost exclusively, supervisory quality risks remain invisible despite its clear relevance to researcher wellbeing, professional development and career continuity. Integrating mentorship more explicitly into funding assessments could therefore help reinforce its importance across the research system.

A practical step would be to assign a modest component of grant evaluation to mentorship quality. For established supervisors, this might include prior mentoring records, ECR outcomes, or institutional evaluation data. For newer supervisors lacking documented history, greater emphasis could be placed on a mentorship plan and evidence of completed supervisory training. Requiring applicants to articulate how they intend to support communication, workload management, inclusion and professional development, would help frame mentorship as an expected dimension of responsible research leadership.

Funding agencies could also strengthen positive incentives by providing additional support to supervisors with consistently strong mentorship records, for example through resources dedicated to team development or training. At the same time, accountability mechanisms are also needed. For grant renewal applications, a proven record of excellence mentorship could be a key non-scientific criterion for favorable consideration. In cases of severe misconduct or persistent supervisory failure, funding bodies could take institutional sanctions into account when assessing eligibility for new grants, particularly those involving personnel support or training responsibilities. Where necessary, previous mentorship scores (or documented institutional ethical violations) could serve as a veto factor, even if the scientific score is high. Requiring institutions to report both sanctions and documented resolution processes could further reduce the risk that serious issues remain without consequences across funding cycles or institutional moves.

Although the implementation of such measures would require coordination across institutions, governments and funding agencies, meaningful improvement in supervisory practices is unlikely without structural commitment at multiple levels. Supportive supervision should therefore be understood not as an individual attribute left to change, but as a core component of research culture that warrants institutional recognition, evaluation and accountability.

## DECLARATIONS

### ETHICS APPROVAL AND CONSENT TO PARTICIPATE

The study received ethical approval from the Ethics Committee for Research Involving Human Participants of the University of the Basque Country (UPV/EHU) (Protocol number: CE PI_2024_013). All participants provided informed consent before accessing the questionnaire. Survey available in the Supplementary Material section.

### AUTHOR CONTRIBUTIONS

All authors contributed equally to the conception and design of the study, as well as to material preparation, data collection, and analysis. The first draft of the manuscript was written by all authors, and all authors commented on previous versions of the manuscript. All authors read and approved the final manuscript.

## ACKNOWLEDGEMENTS

We are deeply grateful to the more than 2,600 individuals from 65 countries who generously took the time to complete our survey and share their experiences. This work would not have been possible without their openness and courage. We also sincerely thank all those who helped disseminate the survey through their platforms, including the social media managers of the Instagram accounts @phddiarymemes and @phdhelp, Dr. Raúl Peña (social media manager of @research_goes_slowly) and the manager of the LinkedIn page “Doing A PhD In Africa.” We are especially thankful to Dr. Andrew Akbashev and Dr. Emmanuel Tsekleves for promoting the survey within their networks and to Dr. Miren Ioar de Guzman and Sonia Olaechea for their ongoing suggestions and support in the dissemination strategy. Lastly, we are grateful to Dr. Leire Escajedo for her valuable legal advice during the development of this project.

## SUPPLEMENTARY FIGURES

**Fig. S1.**
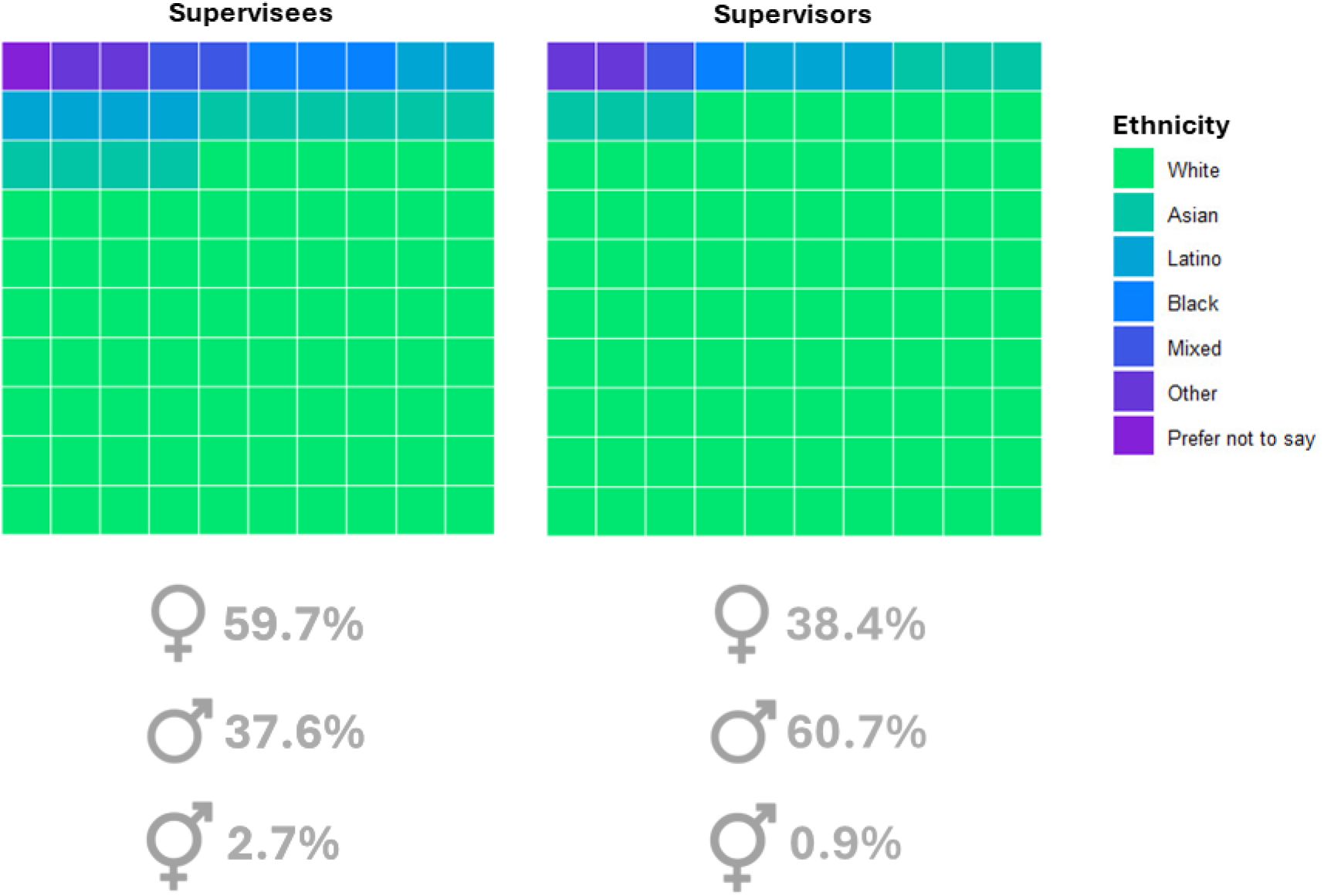
Demographics of respondents and their supervisors. Racial/ethnic composition of both ECRs and supervisors is shown as reported by ECRs. Icons below each panel indicate the gender distribution within each group (female, male, and non-binary/other).

**Fig. S2.**
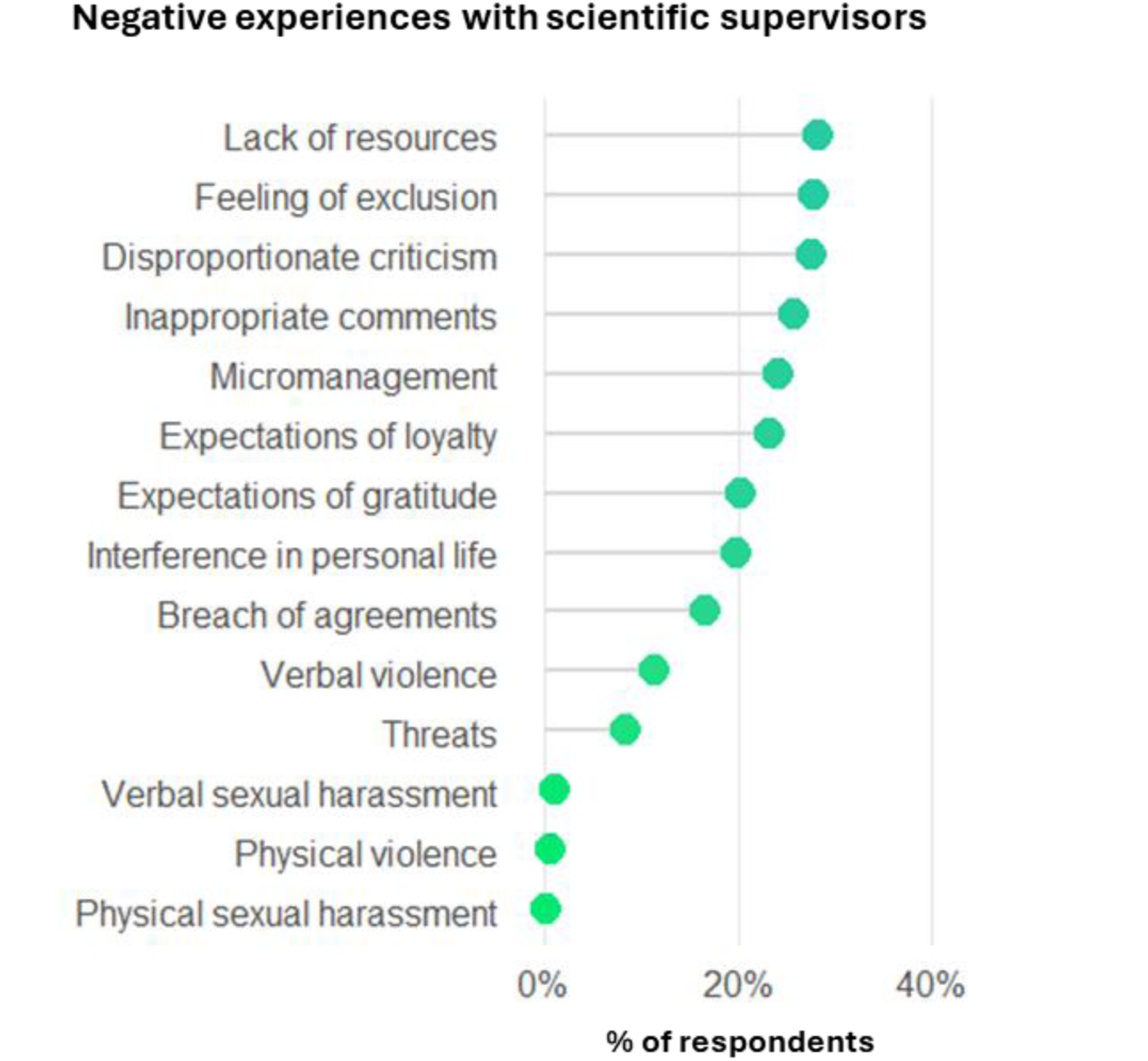
Reported negative experiences with scientific supervisors. Percentage of respondents reporting each situation (ranked 11^th^ and below by overall prevalence; the 10 most frequently reported experiences are shown in Fig. 1).

**Fig. S3.**
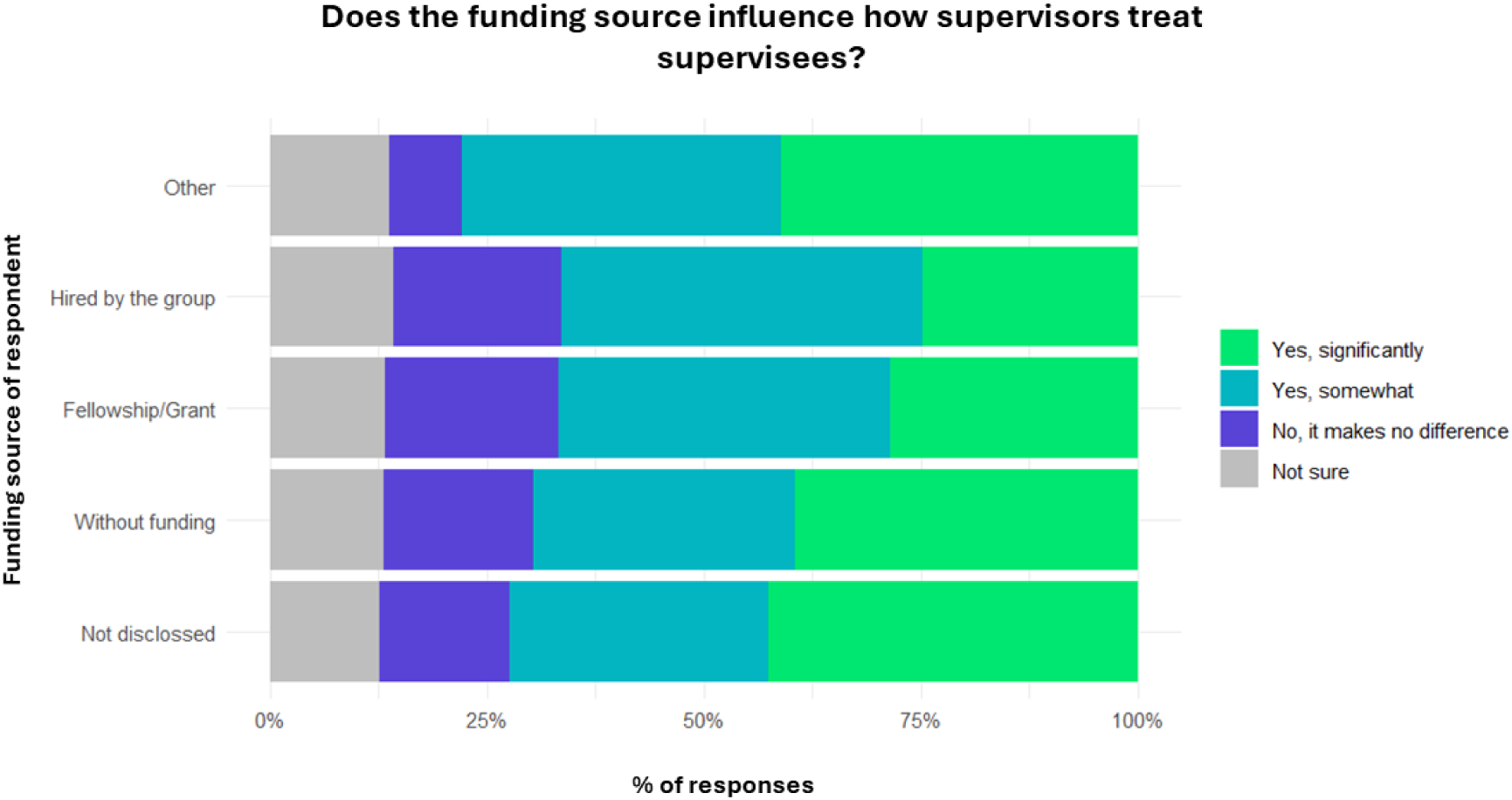
Funding source and supervisory dynamics. Respondent agreement to whether funding status affects ECR-supervisor relationships.

**Fig. S4.**
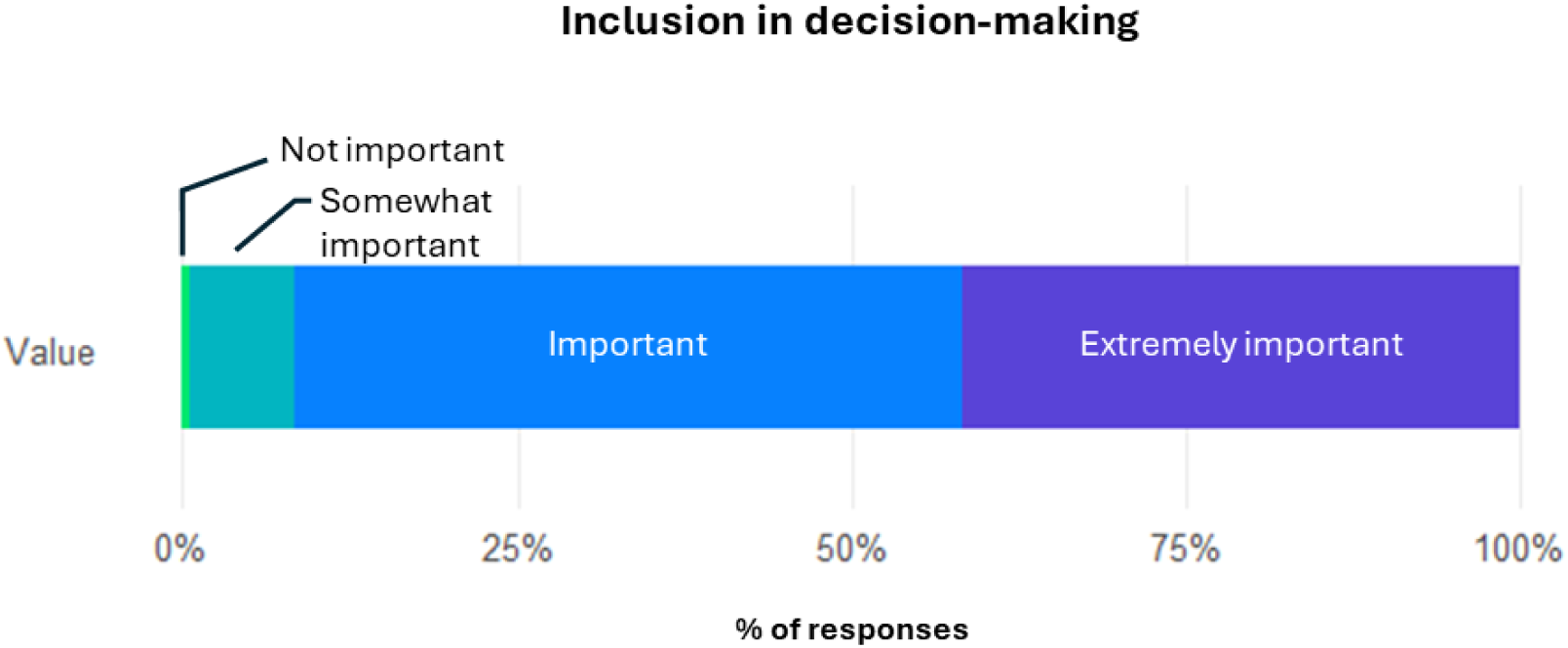
Inclusion of ECRs in decision-making. The bar shows how respondents rated the ability of supervisors to include team members in decision-making.

**Fig. S5.**
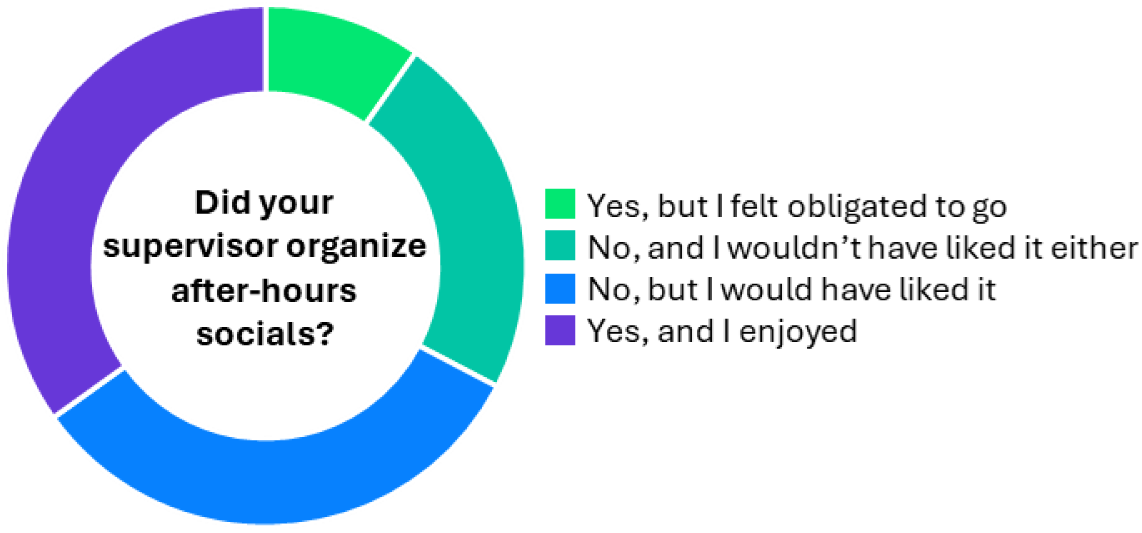
After hours leisure activities. Respondents’ experiences with supervisor-organized after-hours social gatherings.

## SUPPLEMENTARY MATERIAL: SURVEY QUESTIONS

### SECTION 1: DEMOGRAPHIC INFORMATION

**1. What is your current position?**

Predoctoral researcher

Postdoctoral researcher

Former researcher (left academia or went to industry)

**2. If you are a predoctoral researcher, what is your funding status?**

Funded by a fellowship/grant

Hired by the group

Without funding (only student)

**3. If you are a postdoctoral researcher, what is your funding status?**

Funded by a fellowship/grant

Hired by the group

Other

**3.1. If you selected ‘Other’ in the previous answer, please specify here.**

**4. If you are a former academic researcher, what were the main reasons you decided to leave? (choose as many as apply)**

Lack of stable job opportunities

Low salary or financial stability

Work-life balance concerns

High pressure/stressful environment

Lack of funding

Desire to pursue other professional path

Negative experiences with principal investigator or scientific supervisor

Other

**4.1. If you selected ‘Other’ in the previous answer, please specify here.**

**5. What field of research are you involved in?**

Life Sciences

Chemistry Sciences

Physical Sciences

Engineering and Technology

Medical and Health Sciences

Social and Behavioural Sciences

Mathematics and Statistics

Computer and Information Sciences

Humanities and Arts

Bussiness and Economics

Education Sciences

Interdisciplinary Studies

Other

**5.1. If you selected ‘Other’ in the previous answer, please specify here.**

**6. How many years of research experience do you have (including PhD thesis)?**

Less than 1 year

1-3 years

4-6 years

More than 6 years

**7. How many supervisors have you worked with during your research career?**

1

2

3

4 or more

**8. Which country are you currently working in?**

**9. What is your gender?**

Male

Female

Non-binary/Other

Prefer not to say

**10. What is your ethnicity?**

Asian

Black

Latino

Mixed

White

Prefer not to say

Other

**10.1. If you selected ‘Other’ in the previous answer, please specify here.**

**11. What is the gender of your scientific supervisor?**

Male

Female

Non-binary/Other

**12. What is the ethnicity of your supervisor?**

Asian

Black

Latino

Mixed

White

Other

**12.1. If you selected ‘Other’ in the previous answer, please specify here.**

### SECTION 2: CHARACTERISTICS OF GOOD SCIENTIFIC SUPERVISORS

*Please consider the following questions in terms of **<u>general qualities</u> that make a good scientific supervisor**, rather than thinking of any specific individual. Your responses should reflect what you believe are **universally positive traits** for supporting and guiding researchers constructively*.

**1. In your opinion, what are the top <u>three specific</u> characteristics that make a supervisor <u>supportive and constructive</u>?**

Clear project and collaboration management

Balanced workload for team members

Supportive supervisor rather than a boss

Flexibility with time and work hours

Support for the professional development of team members

Regular meetings and communication

Promotion of collaborations among group members

Setting realistic goals and objectives

Maintaining clear records of each member’s contributions in manuscripts/projects

Encouraging experienced postdocs to mentor junior researchers

Respecting personal time and life outside work

Communicating clarity in research risks, benefits, and project expectations

Other

**1.1. If you selected ‘Other’ in the previous answer, please specify here.**

**2. How important do you think regular meetings (with your scientific supervisor) are for effective work management?**

Extremely important

Important

Somewhat important

Not important

**3. How important is the principal investigator’s ability to include team members in decision-making?**

Extremely important

Important

Somewhat important

Not important

**4. What <u>three main strategies</u> do you believe scientific supervisors can take to maintain a balanced workload for team members?**

Having realistic expectations and deadlines

Regularly checking in on workload and progress

Distributing tasks based on individual strengths and interests

Allowing flexibility in work hours and remote work options

Encouraging open communication about workload concerns

Implementing project management tools to track progress

Providing training on time management

Other

**4.1. If you selected ‘Other’ in the previous answer, please specify here.**

**5. How do you think a supervisor’s flexibility with time influences team morale and productivity?**

Significantly improves morale and productivity

Somewhat improves morale and productivity

Has little impact on morale and productivity

Detracts from morale and productivity

Not sure / No opinion

### SECTION 3: CHARACTERISTICS OF BAD SCIENTIFIC SUPERVISORS

*Please consider the following questions in terms of **<u>general qualities</u> that make a bad scientific supervisor**, rather than thinking of any specific individual. Your responses should reflect what you believe are **universally negative traits** that can hinder the growth and well-being of researchers*.

**1. What <u>three main strategies</u> do you believe principal investigators should <u>avoid</u> to prevent an unbalanced workload for team members?**

Having unrealistic expectations and deadlines

Ignoring team members’ workload and progress

Distributing tasks without considering individual strengths and interests

Enforcing rigid work hours without flexibility

Discouraging open communication about workload concerns

Failing to implement project management tools to track progress

Not providing training on time management

Other

**1.1. If you selected ‘Other’ in the previous answer, please specify here.**

**2. What <u>three main actions</u> do you believe scientific supervisors should <u>avoid</u> that could negatively impact team morale and productivity?**

Being inflexible with work hours and remote work options

Micromanaging team members’ work

Providing little or no feedback on performance

Ignoring team members’ personal lives and well-being

Adding authors to your work (manuscripts, conferences, etc.) without consulting you or considering your opinions

Failing to acknowledge achievements of team members

Promoting a competitive environment among team members

Setting unrealistic goals and expectations

Using dismissive or disrespectful communication

Other

**2.1. If you selected ‘Other’ in the previous answer, please specify here.**

**3. What <u>three main practices</u> should supervisors <u>avoid</u> to support their team members’ professional development effectively?**

Failing to offer mentorship and guidance

Not providing opportunities for training and workshops

Discouraging attendance at conferences and seminars

Not promoting collaborations with other researchers

Avoiding collaborations among researchers within the group

Ignoring discussions about career goals and progress

Not offering feedback on research and professional skills

Other

**3.1. If you selected ‘Other’ in the previous answer, please specify here.**

### SECTION 4: PERSONAL EXPERIENCES AND RECOMMENDATIONS

*In this section, for the questions related to supervisors, please **focus your answers on a specific supervisor**, whether current or from the past*.

**1. What was your primary motivation for pursuing a career in academic science?**

Vocation/passion for science

Job stability or financial security

Professional recognition (better positions, prestige)

Other

**1.1. If you selected ‘Other’ in the previous answer, please specify here.**

**2. Did you have any work experience before starting your PhD thesis (outside academia)?**

Yes

No

**3. How would you describe your approach to academic work?**

I mostly work during my scheduled hours and maintain a balance with personal time

I often work beyond my scheduled hours to meet my own or others’ expectations/goals

My work in academia involves constant personal commitment and frequent personal sacrifices

**3.1. Are you satisfied with your current approach?**

Very satisfied and I would not change a thing

Generally satisfied, but with room for improvement

Somewhat dissatisfied, I would like to make changes

Dissatisfied, I feel this approach is not working well for me

**4. Do you believe that being hired directly by the scientific supervisor, as opposed to having your own funding (e.g., fellowships, grants or projects), could influence how the supervisor treats you (e.g., increased pressure or different expectations)?**

Yes, significantly

Yes, somewhat

No, it makes no difference

Not sure

**5. The person you are thinking of was your scientific supervisor during your time as:**

Predoctoral researcher

Early postdoctoral researcher (<4 years)

Senior postdoctoral researcher (>4 years)

**6. Would you say the number of meetings with your scientific supervisor was:**

Too many

Sufficient

Insufficient

**7. Would you describe the usual meetings with your scientific supervisor as:**

Extremely useful

Useful

Useless

A complete waste of time

**8. Did your scientific supervisor organize group leisure activities outside of working hours?**

Yes, and I enjoyed

Yes, but I felt obligated to go

No, but I would have liked it

No, and I wouldn’t have liked it either

**9. Have you experienced any of these situations with your scientific supervisor?**

Abrupt changes in attitude

Interference in personal life

Lack of empathy

Hypocrisy

Disproportionate criticism

Expectations of loyalty

Expectations of gratitude

Feeling of exclusion

Breach of agreements

Lack of resources

Unequal treatment compared to other colleagues

Lack of support

Toxic work environment

Disorganization

Lack of recognition

Micromanagement

Lack of communication

Contact outside of working hours

Assignment of tasks beyond your job responsibilities

Inappropriate comments

Threats

Verbal violence

Physical violence

Verbal sexual harassment

Physical sexual harassment

None of these

**10. To what extent do you think your scientific supervisor’s attitude affected your mental health (positively or negatively)**

Significantly

Moderately

Slightly

None

**11. Have you had a particularly positive or negative experience with a scientific supervisor? If so, please share a brief description. (Please do not include any personal data or information about the scientific supervisor or research center)**

**12. What recommendations would you give to improve the relationship between scientific supervisor and their team members?**

**13. On a scale of 0 to 10, how would you rate the overall value of your supervisor’s role, taking into account both positive and negative aspects of your experience?**

0

1

2

3

4

5

6

7

8

9

10

## REFERENCES

Al Makhamreh, M. & Stockley, D. Mentorship and well-being. Int. J. Mentor. Coach. Educ. 9, 1–20 (2020). 10.1108/IJMCE-02-2019-0013

Busch, C. A., Wiesenthal, N. J., Gin, L. E. & Cooper, K. M. Behind the graduate mental health crisis in science. Nat. Biotechnol. 42, 1749–1753 (2024). 10.1038/s41587-024-02457-z

Cornejo-Araya, C. A., Gallardo-Lazo, M. E., Salas, G., Tapia-Parada, C., López-López, W. & Marsico, G. Well-being in doctoral students: considerations for the academic community. Psychol. Hub 41, 2 (2024). 10.13133/2724-2943/18298

Evans, T. M., Bira, L., Gastelum, J. B., Weiss, L. T. & Vanderford, N. L. Evidence for a mental health crisis in graduate education. Nat. Biotechnol. 36, 282–284 (2018).

Feizi, S. & Elgar, F. Satisfaction, research productivity, and socialization in doctoral students: Do teaching assistantship, research assistantship and the advisory relationship play a role? Heliyon 9, e19332 (2023). 10.1016/j.heliyon.2023.e19332

Gewin, V. Mental health: Under a cloud. Nature 490, 299–301 (2012). 10.1038/nj7419-299a

Hund, A. K., Churchill, A. C., Faist, A. M., Havrilla, C. A., Love Stowell, S. M., McCreery, H. F., Ng, J., Pinzone, C. A. & Scordato, E. S. Transforming mentorship in STEM by training scientists to be better leaders. Ecol. Evol. 8, 9962–9974 (2018). 10.1002/ece3.4527

Jackman, P. C., Jacobs, L., Hawkins, R. M. & Sisson, K. Mental health and psychological wellbeing in the early stages of doctoral study: a systematic review. Eur. J. High. Educ. 12, 293–313 (2022). 10.1080/21568235.2021.1939752

Jones-White, D. R., Soria, K. M., Tower, E. K. & Horner, O. G. Factors associated with anxiety and depression among US doctoral students: Evidence from the gradSERU survey. J. Am. Coll. Health 70, 2433–2444 (2022). 10.1080/07448481.2020.1865975

Lacey, S., Haven, T., Santos, R., Farrelly, T., Murray, M. & Kavouras, P. A roadmap to good practice for training supervisors and leadership: a European perspective. Front. Res. Metr. Anal. 10, 1531467 (2025). 10.3389/frma.2025.1531467

Le, M., Pham, L., Kim, H. & Bui, N. The impacts of supervisor–PhD student relationships on PhD students’ satisfaction: A case study of Vietnamese universities. J. Univ. Teach. Learn. Pract. 18, 18 (2021). 10.53761/1.18.4.18

Levecque, K., Anseel, F., De Beuckelaer, A., Van der Heyden, J. & Gisle, L. Work organization and mental health problems in PhD students. Res. Policy 46, 868–879 (2017). 10.1016/j.respol.2017.02.008

Lovitts, B. E. Leaving the ivory tower: The causes and consequences of departure from doctoral study (Bloomsbury Publishing, 2001).

Maestre, F. T. Ten simple rules towards healthier research labs. PLoS Comput. Biol. 15, e1006914 (2019). 10.1371/journal.pcbi.1006914

National Academies of Sciences, Engineering, and Medicine. The science of effective mentorship in STEMM (National Academies Press, Washington, DC, 2019).

OECD. Adult education level. OECD Data. https://www.oecd.org/en/data/indicators/adult-education-level.html (accessed 27 June 2025).

Pizzolato, D. & Dierickx, K. Research integrity supervision practices and institutional support: A qualitative study. J. Acad. Ethics 21, 427–448 (2023).

Powell, K. Young, talented and fed-up: scientists tell their stories. Nature 538, 446–449 (2016). 10.1038/538446a

Rahal, R. M. et al. Quality research needs good working conditions. Nat. Hum. Behav. 7, 164–167 (2023). 10.1038/s41562-022-01508-2

Rillig, M. C. Ten simple rules for how you can help make your lab a better place as a graduate student or postdoc. PLoS Comput. Biol. 18, e1010673 (2022). 10.1371/journal.pcbi.1010673

Sarabipour, S. et al. Insights from a survey of mentorship experiences by graduate and postdoctoral researchers. bioRxiv (2023). 10.1101/2023.05.05.539640

Sarabipour, S., Macklin, P. & Niemi, N. M. Improving academic mentorship practices. Nat. Hum. Behav. 8, 1228–1231 (2024). 10.1038/s41562-024-01910-y

Solomon, S. & Du Plessis, M. Experiences of precarious work within higher education institutions: A qualitative evidence synthesis. Front. Educ. 8, 960649 (2023).

Tenenbaum, H. R., Crosby, F. J. & Gliner, M. D. Mentoring relationships in graduate school. J. Vocat. Behav. 59, 326–341 (2001). 10.1006/jvbe.2001.1804

Tenorio-Lopes, L. Mentor-supervisee relationships in academia: insights toward a fulfilling career. Front. Educ. 8, 1198094 (2023). 10.3389/feduc.2023.1198094

Vigil-Avilés, D. J., Jang, Y. & Urban, M. ‘Take a break, you’ll be able to work more’: convergent mixed methods analysis of PhD students’ blog posts. Stud. Contin. Educ. (2024). 10.1080/0158037x.2024.2319806

Williams, S. N., Thakore, B. K. & McGee, R. Coaching to augment mentoring to achieve faculty diversity: a randomized controlled trial. Acad. Med. 91, 1128–1135 (2016). 10.1097/ACM.0000000000001026

Woolston, C. Paths less travelled. Nature 562, 611–614 (2018). 10.1038/d41586-018-07111-8

